# High-frequency axonal bursts mediate bidirectional modulation of dopamine signaling by nicotinic receptors

**DOI:** 10.64898/2025.12.19.695584

**Authors:** Paul F. Kramer, Anthony Yanez, Faye Clever, Renshu Zhang, Zayd M. Khaliq

**Affiliations:** Cellular Neurophysiology Section, National Institute of Neurological Disorders and Stroke, National Institutes of Health, Bethesda, MD, 20892, USA; Aligning Science Across Parkinson’s (ASAP) Collaborative Research Network, Chevy Chase, MD 20815, USA; Department of Molecular, Cellular, and Developmental Biology, University of Michigan, Ann Arbor, MI, 48109, USA; Biochemistry, Cellular and Molecular Biology Graduate Program, Johns Hopkins School of Medicine, Baltimore, MD, 21287, USA; Department of Molecular & Cellular Biology, University of California, Berkeley, Berkeley, CA, 94720, USA

## Abstract

Nicotinic acetylcholine receptors (nAChRs) facilitate striatal dopamine transmission but also suppress dopamine release during high-frequency stimulation, suggesting they act as as low pass filters of dopamine release. Because axonal excitability is key a determinant of transmission, we combined axonal recordings and calcium imaging to define the physiological conditions under which nAChRs bidirectionally control dopaminergic axons. Activation of cholinergic interneuron (CINs) recruited nAChRs to enhance dopaminergic axon signals under moderate activation but suppressed signals after strong high-frequency stimulation. Axonal recordings revealed that single-pulse striatal stimulation triggered a rapid (∼125 Hz) burst of 2-3 nAChR-driven spikes in dopaminergic axons followed by a brief refractory period that inhibited further axon spiking. In sum, we show that nAChRs mainly enhance local excitability of striatal dopaminergic axons but also trigger axonal bursting that suppresses axonal excitability. This mechanism expands the computational power of dopaminergic axons and explains the apparent nAChR-mediated low-pass filtering of dopamine release.

## INTRODUCTION

Dopaminergic neurons of the ventral midbrain (DANs) regulate behaviors such as motivation, reward learning, and saliency through striatal dopamine (DA) release (Berke, 2018; Costa & Schoenbaum, 2022; Gershman et al., 2024; Wise & Jordan, 2021). Signaling in DANs results from the interaction of synaptic inputs in the soma and dendrites that regulate the generation of action potentials (APs) at the axon initial segment (Hausser et al., 1995). These APs propagate to terminals, triggering DA release, thus relaying somatodendritic activity. In addition to this standard view of neuronal signaling, dopaminergic axons express receptors that are positioned to locally modulate striatal DA release in a manner that is distinct from somatodendritic signaling (Seddon & Kramer, 2025). Local receptor modulation of dopaminergic axonal excitability represents a distinct strategy to shape signaling from DANs onto targets within the striatum.

The axons of DANs express nicotinic acetylcholine receptor (nAChR), activated by transmission from striatal cholinergic interneurons (CINs) (Brimblecombe et al., 2018). However, the effects of axonal nAChRs on DA signaling are still unclear. There is evidence that nAChRs promote DA release, which is involved in movement and motivation in mice (Liu et al., 2022; Mohebi et al., 2019) and learning in song birds (Qi et al., 2025). Other evidence suggests no causal relationship in mice (Azcorra et al., 2023; Chantranupong et al., 2023; Krok et al., 2023), or perhaps that nAChRs play an opposing role where they inhibit striatal DA release (Zhang et al., 2025). These data highlight the need for better mechanistic understanding at the cellular level.

Physiological studies have highlighted both excitatory and inhibitory roles for axonal nAChRs in modulating dopamine release, but the mechanisms distinguishing these roles remain unresolved. Optical or spontaneous activation of striatal CINs robustly drives nAChR-mediated DA release in both dorsal and ventral striatal regions (Cachope et al., 2012; Matityahu et al., 2023; Threlfell et al., 2012; Yorgason et al., 2017; Zhou et al., 2001). The mechanism driving CIN-mediated DA release is nAChR-mediated initiation of APs in the dopaminergic axon (Liu et al., 2022) through a novel form of neurotransmission between axons (Kramer et al., 2022). Conversely, early studies suggested an additional role of nAChRs in inhibiting DA release. Activation of nAChRs potentiated DA release for bursts of stimulations at low-frequencies (<20 Hz), but suppressed DA release following high-frequency (>20 Hz) stimulations (Rice & Cragg, 2004; Zhang & Sulzer, 2004), suggesting that axonal nAChRs function as low-pass filters that amplify low-frequency DA release but dampen DA release at high-frequencies. The mechanisms underlying low-pass filtering and the nAChR-dependent suppression of signaling in dopaminergic axons has not be determined.

Here, we examined how nAChRs may bidirectionally modulate (ie. enhance or suppress) the excitability of dopaminergic axons using a combination of patch-clamp voltage measurements and calcium imaging from dopaminergic axons in the dorsomedial striatum. We demonstrate that activation of axonal nAChRs consistently produced subthreshold depolarization and AP firing, confirming their role as excitatory receptors. Calcium imaging experiments examining coincident activation of CINs and dopaminergic axons showed that submaximal CIN activation enhanced dopaminergic axonal excitability. By contrast, strong stimulation of CINs suppresses axonal dopaminergic signals, suggesting that the strength of nicotinic receptor activation modulates DA axon activity. Interestingly, axonal patch-clamp experiments showed that a single electrical shock or light pulse activation of CINs resulted in a high-frequency burst of APs in dopaminergic axons, which we determined are nAChR-dependent. Lastly, we tested axonal excitability following nAChR-mediated bursts and found that bursting induced a period of reduced axonal excitability lasting 60-80 ms. These data demonstrate that nAChRs are fundamentally excitatory in DA axons. They also show that strong cholinergic drive can suppress stimulated train-mediated DA release via burst-induced reductions in axonal excitability. This effect produces an apparent low-pass filtering effect when recording dopamine release. Altogether, these findings further support the idea that dopaminergic axons act as dynamic integrators of striatal network activity rather than simple signal propagators, expanding the computational repertoire of these axons to include the initiation of high-frequency bursting.

## RESULTS

### Quantifying nAChR modulation of bursting in DAergic axons by imaging jGCaMP8f

Nicotinic receptors (nAChRs) expressed on the axons of DANs have been suggested to perform a filtering action on striatal DA release, enhancing DA release during single and low frequency stimulation, and suppressing DA release evoked by high-frequency stimulation (Rice & Cragg, 2004; Zhang & Sulzer, 2004). The influence of nAChRs on striatal DA has been primarily examined using carbon fiber voltametric measurements (Adrover et al., 2020; Stouffer et al., 2015; Threlfell et al., 2012; Zhou et al., 2003), which quantify the concentration of DA released from thousands of vesicle release sites (Robinson et al., 2003). However, transmitter release is largely determined by processes upstream of vesicular release, like the frequency and characteristics of axonal AP firing. Whether nAChR-mediated filtering results from effects on axonal excitability is unclear. We tested the effect of nAChR signaling on dopaminergic axonal excitability using electrical stimulation of the dorsomedial striatum in acutely prepared brain sections. This results in co-stimulation of dopaminergic axons by two temporally separated processes: immediate direct excitation from the electrical stimulation and delayed synaptic excitation through acetylcholine (ACh) release from CIN terminals onto axonal nAChRs. We monitored calcium (Ca2+) influx in striatal dopaminergic axons by expressing the genetically-encoded indicator jGCaMP8f (Zhang et al., 2023) in DANs of the SNc (Figure 1A-C). Tonic and high-frequency bursting are characteristic patterns of DANs (Grace & Bunney, 1984). Therefore, we mimicked this activity by stimulating trains of 10 pulses at 2 Hz to produce a tonic pattern of activity, followed by a 750 ms pause and then a burst stimulation of 5 pulses at 50 Hz (Figure 1D).

**Figure 1.**
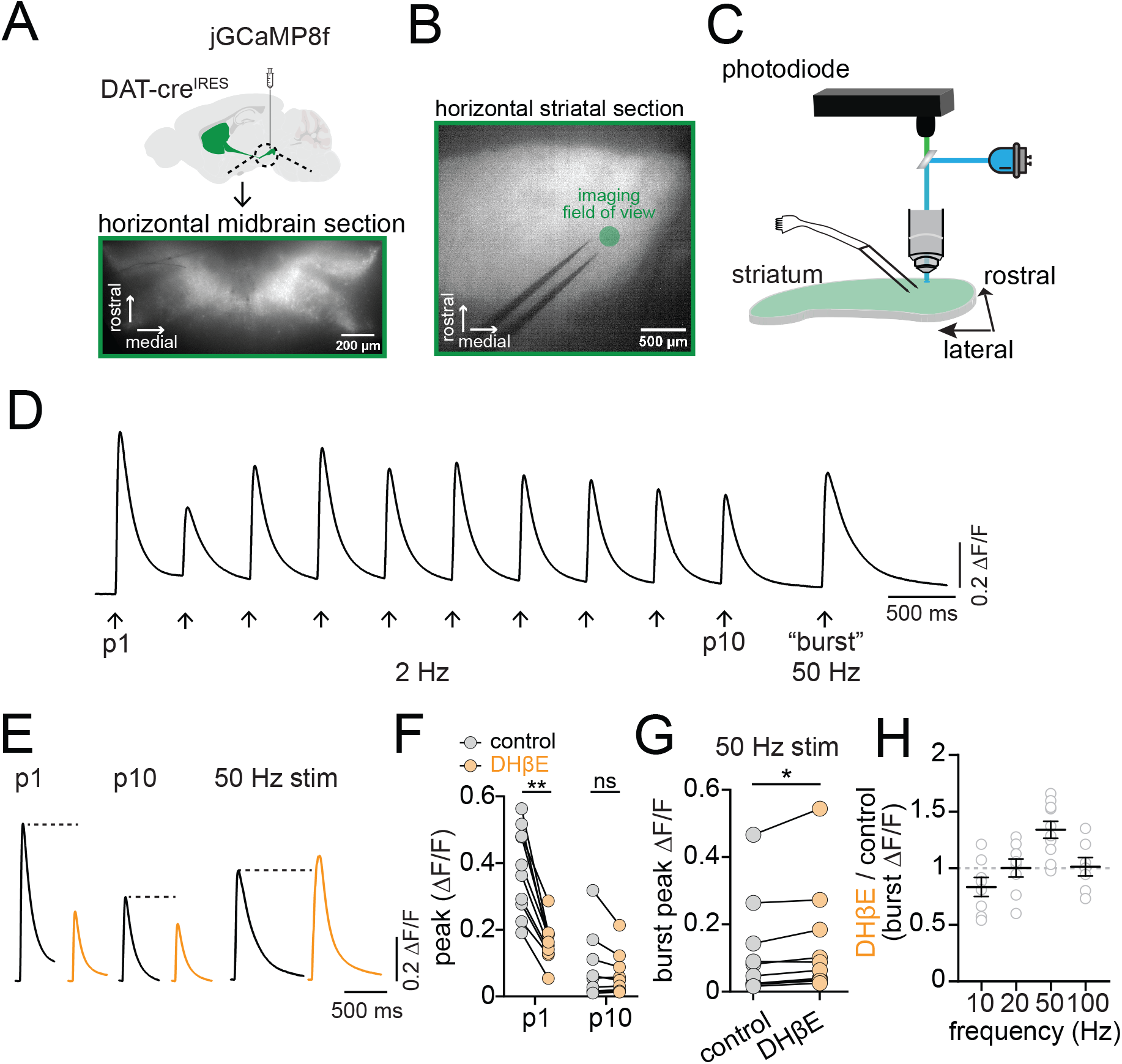
nAChRs inhibit burst-mediated excitability of dopaminergic axons. **A.** Cre-dependent jGCaMP8f AAV injected into the ventral midbrain of DAT-cre mice; lower panel shows infected neurons. **B.** Fluorescent image of horizontal DMS section with jGCaMP8f expression, bipolar electrode, and imaging site (green circle). **C.** Cartoon of experimental setup: LED excitation and photodiode recording of jGCaMP8f fluorescence. **D.** Example photodiode trace showing jGCaMP8f responses to repeated DMS electrical stimulation with 10 stimuli at 2Hz followed by 5 stimuli at 50 Hz. **E.** Example jGCaMP8f traces showing first (P1), last (P10) in 2 Hz train, and 50 Hz burst stimulation in control (black) and DHβE (orange) ACSF. **F.** Collected peak jGCaMP8f signal amplitudes in control (gray symbols) and DHβE (orange symbols) for P1 and P10. **G.** Collected peak signal amplitudes for 50 Hz burst stimulation in control solutions versus DHβE. **H.** Ratio of jGCaMP8f signal amplitudes measured in DHβE over control aCSF for 10, 20, 50 and 100 Hz stimulation frequencies. *p<0.05, **p<0.01, ns > 0.05. All experiments in D2R, GABA-B, and mAChR antagonists.

To examine the contribution of nAChR signaling to dopaminergic axon excitability, we tested the effect of the nAChR antagonist dihydro-p-erythroidine hydrobromide (DHβE, 1 μM) on stimulated dopaminergic axon signals. We found that the amplitude of the first pulse (P1) in the tonic stimulation train was significantly reduced following DHβE inhibition of nAChRs (avg ΔF/F, control=0.38; DHβE=0.16, t(18)=9.7, p<0.0001; Figure 1E and 1F), but the tenth pulse (P10) amplitude was unaffected (avg ΔF/F, control=0.08; DHβE=0.06, t(18)=0.77, p=0.45). By contrast, 50 Hz burst stimulation resulted in a Ca2+ signal that was significantly increased in amplitude following DHβE inhibition of nAChRs (avg burst peak ΔF/F, control=0.12, DHβE=0.14, t(9)=2.95, p=0.016; Figure 1G). In separate experiments, we tested the effect of DHβE on Ca2+ signals following 5 pulse stimulations delivered at 10, 20 and 100 Hz (Figure 1H). Plots of the ratio of Ca2+ signals in DHβE relative to control showed that nAChR inhibition reduced the amplitude of 10 Hz stimulation signals, enhanced the 50 Hz signals and had no effect on signals evoked by 20 Hz and 100 Hz (avg DHβE/control Ca2+ signal, 10 Hz, 0.83 +/- 0.08; 20 Hz, 1.00+/0.08; 100 Hz, 1.01+/-0.08). Therefore, these results demonstrate that nAChRs enhance dopaminergic axon excitability during CIN/DAN axon costimulation at single or low-frequencies below 20 Hz. However, for higher frequency stimulation at 50 Hz, application of nAChR antagonists led to a significant increase in dopaminergic axon Ca2+ signals suggesting they play a complicated, but net inhibitory, role in regulating dopaminergic axon excitability following high-frequency burst stimulation.

### Action potential bursting in distal dopaminergic axons

To determine how nAChRs enhance and inhibit dopaminergic axon excitability at low and high frequencies, respectively, we next recorded the membrane potential of dopaminergic axons in the dorsomedial striatum following single- or burst-stimulation protocols. We found that a single pulse of electrical stimulation could evoke an AP in the dopaminergic axon, as expected. Surprisingly, we also observed that a single stimulus resulted in a high-frequency burst of 2-3 axonal APs (Figure 2A-2C; Supplemental Figure 1A,B). In trials with two or more APs, the first AP occurred 1.74 ± 0.84 ms after the electrical pulse, and the second AP occurred an average of 9.22 ± 1.36 ms after the stimulus (median instantaneous rate, 122 ± 26.2 Hz, n=16). Importantly, we found that all APs occurring more than 6 ms after the stimulus were abolished by DHβE, whereas those occurring earlier were typically unaffected (Figure 2D). Of the 50 trials we analyzed in which nAChR-mediated APs were evoked, we observed a single AP in 38 trials, a doublet in 11, and a triplet in 1 (Figure 2E). Based on these observations, we concluded that the earliest APs resulted from direct excitation of the dopaminergic axon (“direct”), while delayed APs (> 6ms delay from stimulation) were likely evoked by activation of nAChRs.

**Figure 2:**
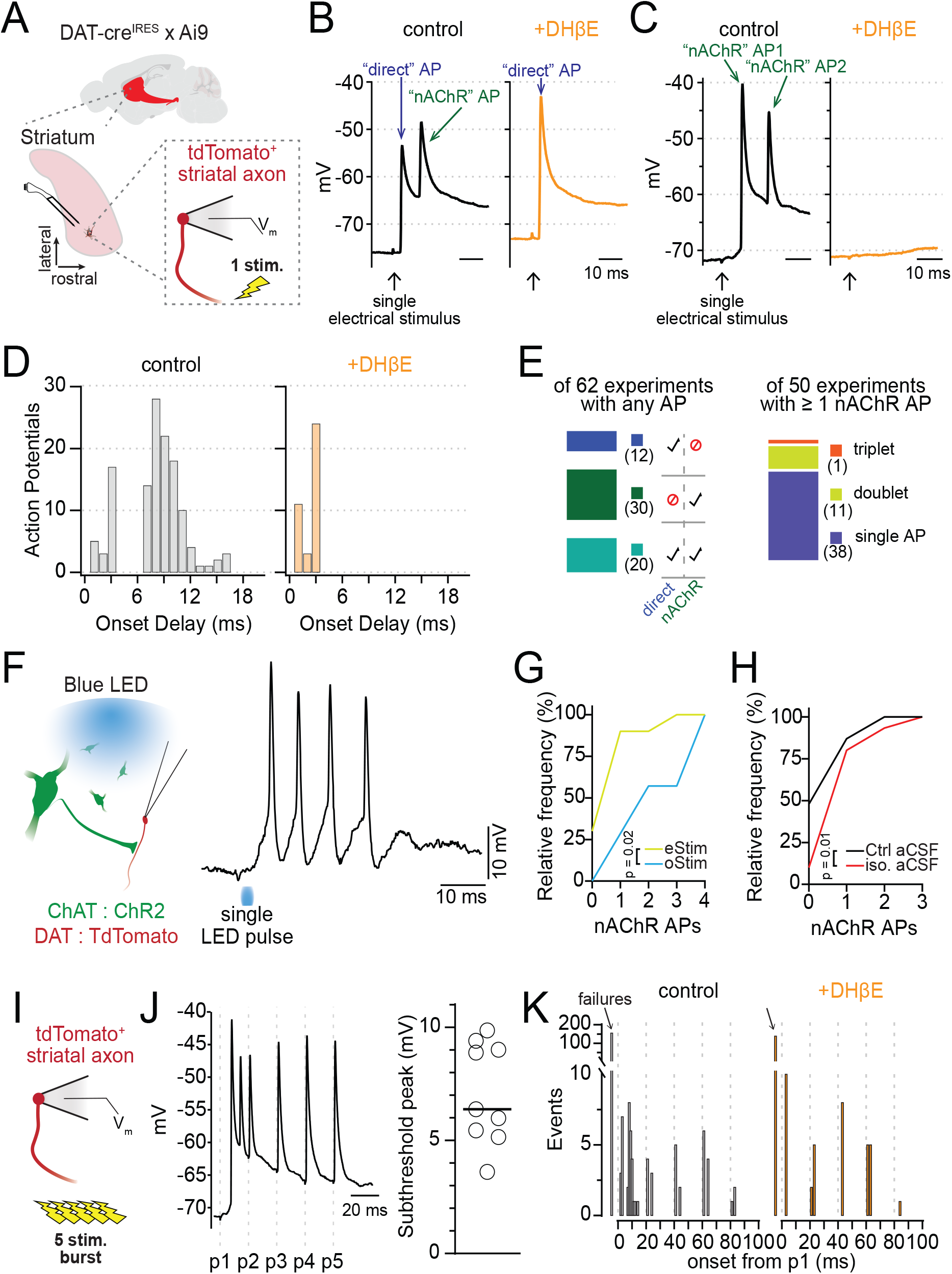
Axonal nAChR signaling initiates action potential bursting. **A.** Diagram of perforated patch recordings from DAT+ axons. B-C. Example traces of two AP burst patterns evoked by single electrical stimuli: direct AP with a nAChR-dependent AP (B), or two nAChR-dependent APs (C). DHβE (1 JM) is a nAChR antagonist. **D.** Frequency distribution of AP onset times in control (left) and DHβE (orange). **E.** (Left) Number of axons recording experiments (n= 62 total) with direct, nAChR-evoked, or both AP types; (Right) subset showing cases with 1-3 nAChR-mediated APs per experiment. **F.** (Left) ChR2 and TdTomato expression in ChAT+ and DAT+ neurons, respectively, via AAV in ChAT-cre × DAT-cre mice. (Right) Example of AP bursts in a DAergic axon following a single blue light pulse in isolation aCSF. **G.** Cumulative frequency: number of experiments with 1–4 APs, compared between electrical and optical stimulation. **H.** Cumulative frequency: number of experiments with 1–3 APs, compared for electrical stimulus experiments in control and isolation aCSF. **I.** Experiment diagram for burst stimulation of DAergic axons. **J.** Example DAergic axon voltage trace during 5-stim, 50 Hz bursts in iso. aCSF and collected peak depolarization of the subthreshold burst envelope. **K.** Frequency distribution of AP onset times during 50 Hz bursts in isolation aCSF (black) and DHβE (orange). Iso. aCSF contained D2R, GABA-B, and mAChR antagonists.

Occasionally, the first AP of the burst was delayed, occurring 9.51 ± 1.54 ms after the stimulation, with the second nAChR-evoked AP at 17.5 ± 2.5 ms. In one case, a third nAChR-evoked AP occurred 20.4 ms following the single electrical pulse. In these cases, the burst frequency was not significantly different from bursts starting with early direct APs (median instantaneous rate, 125 ± 42.8 Hz, p=0.35, n=12; Supplemental Figure 1B). Of note, we also recorded a single example of a spontaneously occurring high-frequency burst of APs (Supplemental Figure 1E), indicating that nAChR-mediated AP bursting in dopaminergic axons is not dependent on external stimulation.

To test the involvement of CINs in nAChR-mediated bursting in dopaminergic axons, we expressed channelrhodopsin (ChR2) in striatal CINs to selectively evoke ACh release from these neurons while recording the membrane voltage from nearby dopaminergic axons (Figure 2F). Similarly, we found that single light pulses evoked robust, high-frequency bursts of 2-4 APs in dopaminergic axons (Figure 2F; Supplemental Figure 1C-D). Comparisons of electrical and optical stimulation showed that optical stimulation evoked a bursting at a higher probability and evoked bursts with greater numbers of APs on average (Figure 2G; D=0.59, p=0.02, n(eStim) = 10, n(oStim) = 7).

The striatum is a dense hub of local interneurons and long-range inputs, many of which alter both the activity of CINs and the terminal release of DA (Burke et al., 2017; Liu & Kaeser, 2019; Ratna & Francis, 2025; Sulzer et al., 2016). Central to the axo-axonic transmission of ACh from CINs to dopaminergic axons are the modulatory neurotransmitters DA, GABA and ACh. To investigate if inhibition from these transmitters alters the initiation of APs in dopaminergic axons, we compared the frequency of nAChR-evoked APs in the presence of inhibitors against muscarinic-, D2-, and GABAb-receptor (“iso. aCSF”, see methods). We observed a significant increase in multispike bursts when recordings were made in iso. aCSF. (p=0.01, control n=46, iso. aCSF n=30; Figure 2H), suggesting that inhibitory GPCRs limit, but do not prohibit, nAChR-dependent bursting in dopaminergic axons. GABA-A receptors can modulate the integration of nAChR activity in dopaminergic axons, but they do so equally for single- or burst-stimulated activity (Brill-Weil et al., 2025).

To determine whether a train of electrical stimulation could evoke more nAChR-mediated spiking, we next evoked a 5-pulse train at 50 Hz with the bipolar electrode while recording voltage from dopaminergic axons of the striatum (Figure 2I). As earlier, we observed a burst of nAChR-mediated APs following the first stimulation and directly stimulated APs in the dopaminergic axons, as well as a broad depolarization envelope of 7.08 +/- 2.24 mV (Figure 2J). Having already established the distinction between direct and nAChR-mediated APs in dopaminergic axons, we subsequently used the timing of these events to classify them. Action potentials with an onset before 3 ms following an electrical stimulation were classified as direct APs, while those with an onset between 7 and 20 ms after an electrical stimulation were classified as nAChR-mediated APs. Using this classification, we observed that nAChR-mediated APs only occurred after the first electrical pulse in the 5-pulse burst (Figure 2K). Pulses 2 through 5 of the train were made up only of direct APs. Supporting this conclusion, the application of DHβE to block nAChR signaling eliminated only those APs with an onset between 7 and 20 ms after the first electrical stimulation (Figure 2K).

In summary, we find that single electrical or optical stimulations of ACh release in brain tissue can evoke AP bursting in dopaminergic axons at very high instantaneous frequencies. Furthermore, we show that optical stimulation of CIN is significantly more effective in driving axonal bursting in dopaminergic axons relative to electrical stimulation.

### Phasic subthreshold depolarization in dopaminergic axons is excitatory

One possible mechanism for the nAChR inhibition of burst-evoked DA release could emerge from the nAChR-mediated axon depolarizations, which may increase sodium channel inactivation and reduce axonal spiking (Zhang et al., 2025). A similar mechanism was proposed to reduce axonal excitability following activation of axonal GABA-A receptors [9]. However, nAChR-mediated membrane depolarization may also favor axonal AP formation [24, 25]. To distinguish these two possibilities, we tested the effect of ACh on evoked firing in dopaminergic axons (Figures 3A and 3B). To assess spiking responses, we obtained perforated-patch recordings from dopaminergic axons in the medial fiber bundle (MFB) and evoked APs using a 50 Hz train of 10 current steps (1 ms duration) of increasing amplitude (+5 pA per sweep). We performed these experiments first under control conditions and then following a puff of 300 μM ACh (Figure 3C).

**Figure 3:**
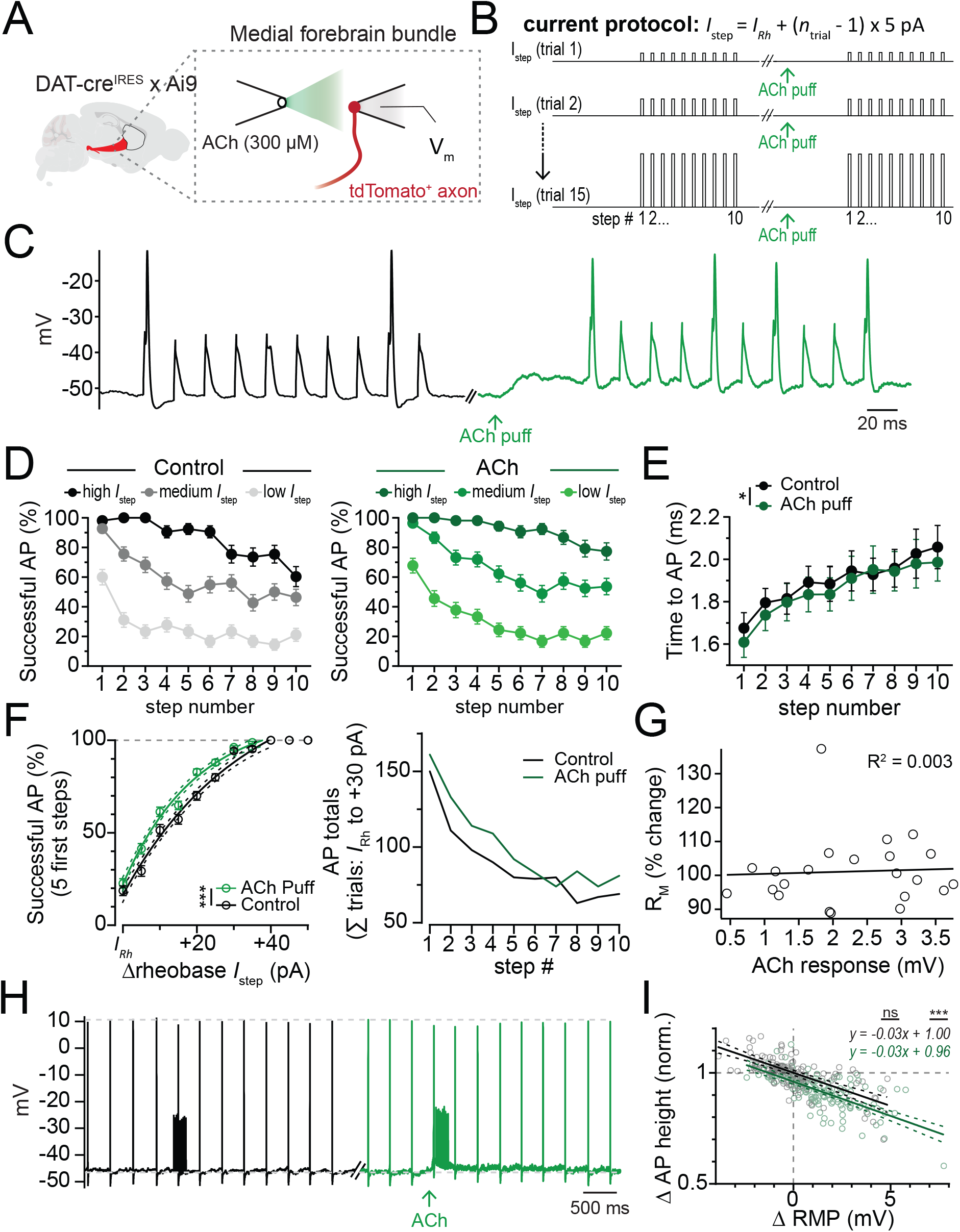
Subthreshold nAChR depolarizations are excitatory. **A.** Experimental setup: DAergic axons labeled via DAT-cre * Ai9 mice; perforated patch recordings from MFB axons; nAChRs activated by local ACh ejection. **B.** Current injection protocol to assess AP initiation probability; trial 1 current (*I*_Rh_) near rheobase. **C.** Example DAergic axon traces during 10-pulse, 50 Hz current trains in control (black) and after ACh pressure ejection (green). **D.** Current injections grouped by amplitude: low, medium, high for control (black) and ACh pressure ejection (green). **E.** Action potential latency per pulse in control and during ACh ejection. **F.** (Left) Percentage of APs generated within the first 5 steps of a 10-step burst for varying current levels near rheobase (X); within-trial comparison in control (black) and ACh ejection (green). (Right) Total APs summed across all current injection levels from 15 axons, compared within trials (control vs. ACh ejection) for current step amplitudes from *I*_Rh_ to *I*_Rh_ + 30 pA. **G.** Change in input resistance during ACh ejection versus nAChR depolarization amplitude. **H.** Example spontaneously active DAergic axon trace showing the change in AP height during control current injection train (black) and for a train in combination with ACh pressure ejection (green). **I.** Change in AP height from changes in axonal resting membrane potential (RMP) for control and ACh pressure ejection. *p<0.05, **p<0.01, ***p<0.001.

We found that puffing ACh resulted in a steady depolarization of the average axonal membrane voltage from -58.25 ± 1.53 mV to -55.69 ± 1.78 mV (n=15), which reversed near 0 mV, as expected for neuronal nAChRs (Vernino et al., 1992) (Supplemental Figure 2). To analyze the effect of this on axonal spiking, we grouped the data into 3 categories based on the strength of the injected current relative to the rheobase current for each recorded axon (IRh; avg IRh = 135 +/- 21.5 pA) - low (IRh to IRh +10 pA), medium (IRh +15 to IRh +25 pA), and high (above IRh +30 pA) current injection groups. We found a significant reduction in the likelihood of AP firing later in the burst train than at the start for all groups (F(9)=18.9, p<0.0001; Figure 3D, left). This effect was similar during nAChR depolarization (F(9)=20.29, p<0.0001; Figure 3D, right). The reduction in axonal excitability over the course of the burst could arise from an increase in potassium conductance, a decrease in sodium channel availability, or some combination. If sodium channel availability decreases throughout the burst, then the time to AP onset (peak of the first AP derivative) should increase concurrently. Consistent with this, the time to onset increased throughout the train from 1.68 ± 0.07 ms on pulse 1 to 2.1 ± 0.1 ms by pulse 10. This effect was similar during nAChR depolarization. Interestingly, the average time to peak was slightly shorter with nAChR activation suggesting an overall increase in excitability (ctrl: 1.90 ± 0.04 ms, ACh: 1.86 ± 0.04 ms, main effect of ACh: F(1,10)=5.45, p=0.04, n=11).

We next examined whether nAChR activation affected the likelihood of evoking an axonal AP. Because there is an intrinsic reduction in axonal excitability over the course of a burst, we examined successes/failures in control or during nAChR activation for the first 5 steps in the train. We found a significantly increased likelihood of AP initiation for all current steps from IRh to +50 pA (main effect of ACh: F(1, 428) = 11.95, p=0.0006; Figure 3F, left). Because the current step amplitude and numbers were consistent between control and nAChR activation, we also counted the total number of action potentials for each current step for step amplitudes between IRh and +30 pA. We found that nAChR activation increased the number of APs throughout the train (control: 887 APs, ACh: 1005 APs out of 1930 current steps, Figure 3F, right).

We next analyzed the relationship between resting membrane potential and axonal spike height (Figure 3H and 3I). We found that activation of nAChRs had no effect on the fitted linear relationship between axonal spike height and resting membrane potential (slope, control, -0.031; ACh:-0.029; F(1, 278)=0.05, p=0.82). However, the Y-intercept was lower with nAChR activation than in control (y-intercept, control: 1.0, ACh: 0.96; F(1,278)=23.6, p<0.0001). Combined with the earlier result showing no detectable change in the axonal input resistance following nAChR activation (Figure 3G), this result suggests a subthreshold nAChR depolarization reduces spike peak, likely from sodium channel inactivation (Figure 3H and 3I).

Together, these results show that nAChR-mediated depolarizations increase the likelihood of local AP initiation. In addition, dopaminergic axons become less excitable during a 50 Hz AP train. Lastly, nAChR depolarization leads to a small but measurable decrease in the amplitude of propagating APs, which likely results from sodium channel inactivation rather than shunting.

### nAChRs suppress dopaminergic axonal excitability as a consequence of robust excitation

*In vivo*, CIN activity within the local striatal network gives rise to varying levels of ACh release from tonic or burst activity (Aosaki et al., 1995; Wilson et al., 1990), resulting in a range of nAChR activation on dopaminergic axons. To test the sensitivity of Ca2+ responses in dopaminergic axons across a range of nAChR activation intensities, we imaged axonal Ca2+ signals while systematically varying electrical stimulation strength and constructed stimulation-response curves (see methods). We then repeated these experiments in the presence of DHβE to test the contribution of nAChRs to Ca2+ signals at different stimulation intensities. We found that single stimulus-evoked Ca2+ signals were reduced across all stimulation intensities by DHβE application (control-DHβE peak dF/F, stim 1x, t(8)= 3.8, p=0.02; stim 3x, t(8)= 4.9, p=0.005; stim 10x, t(8)=4.6, p=0.007; stim 30x, t(8)=4.5, p=0.008), consistent with previous reports that nAChRs potentiate DA release (Figure 4A and 4B). Similarly, burst-evoked Ca2+ signals in response to low-intensity stimulation were significantly reduced by DHβE. However, the burst responses to strong stimulation were unaffected by DHβE (Figure 4A and 4C; control-DHβE peak dF/F, stim 1x, t(8)=4.8, p=0.006; stim 3x, t(8)=4.3, p=0.01; stim 10x, t(8)=2.5, p=0.14; stim 30x, t(8)=1.6, p=0.47).

**Figure 4.**
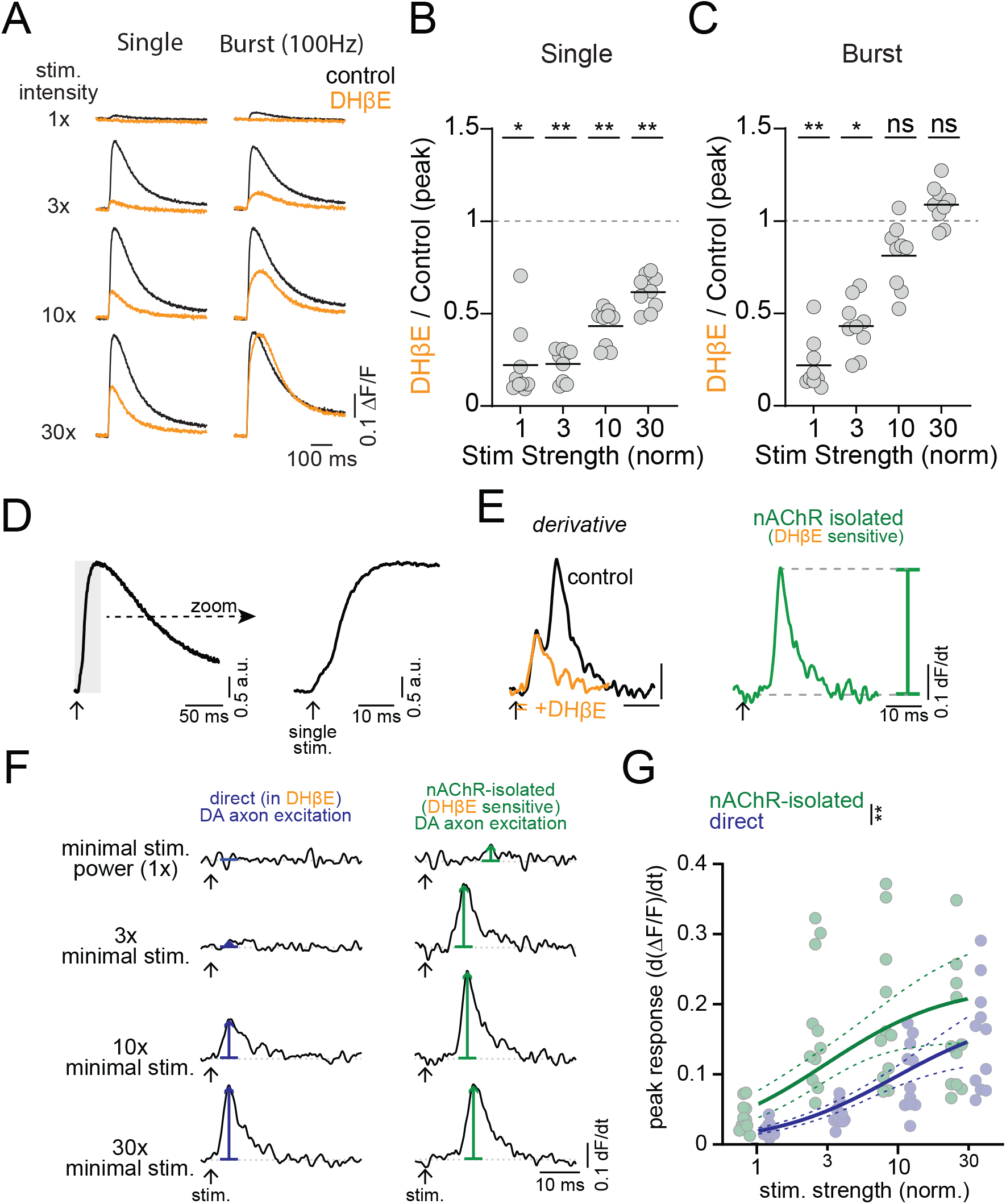
Weak stimulations produce nAChR excitation while strong stimulations produce nAChR inhibition. **A.** Example traces for single and burst (5 stim at 100 Hz) protocols in control aCSF (black) and after nAChR inhibition with DHβE (1 μM, orange), across stimulation intensities. **B.** Inhibition of single-stimulus peak jGCaMP response by DHβE as a function of intensity. **C.** Inhibition of burst peak jGCaMP response by DHβE as a function of intensity. N=9 slices D-E. Analysis workflow for isolating direct and nAChR-mediated components from compound jGCaMP8f photodiode signals. **F.** Example differentiated jGCaMP8f traces showing direct DAergic axon activation (left, blue) and nAChR-mediated activation (right, green) across increasing stimulation intensities (top = minimal intensity). **G.** Peak differentiated jGCaMP8f signals for direct (blue) and nAChR-mediated (green) excitation plotted against normalized stimulation strength. Dashed lines on curve fits are 95% confidence bands. *p<0.05, **p<0.01, ns: p>0.05. Experiments performed with D2R, GABA-B, and mAChR antagonists.

Our results suggest that the interaction between CINs and nAChRs on DA axons depends on the intensity of the cholinergic stimulus, particularly during burst stimuli. To better understand this interaction, we distinguished Ca2+ influx due to nAChR excitation from that due to direct dopaminergic axon stimulation by differentiating the axonal Ca2+ signals (Brill-Weil et al., 2025)(Figure 4D and 4E). The differentiated Ca2+ signal was biphasic, consisting of an early peak followed by a second, larger peak that was delayed in its timing. Following application of DHβE, the late second peak was abolished, consistent with the conclusion that it originates from nAChR activation (Brill-Weil et al., 2025). Plotting the differentiated Ca2+ signals for nAChR and direct dopaminergic axon evoked components against stimulation strength, we found that the half maximal response of the nAChR excitation occurred at a lower intensity than for direct dopaminergic axon stimulation (Kd F-test: 1,84) = 9.1; p = 0.003; Figure 4F and 4G). This observation is consistent with our findings of the undifferentiated signals, which show that nAChR-evoked signals dominate at weak to moderate stimulation strengths. Thus, CIN-mediated Ca2+ signals in dopaminergic axons are more sensitive to stimulation than the direct dopaminergic axon component, supporting the idea that dopaminergic axons are primarily excited by nAChRs over a broad range of activation intensities. Burst-evoked DA release is most prominent at the lowest stimulation intensities, whereas with increasing stimulation strength, nAChRs lead to little further effect on dopaminergic axonal excitability or may even suppress responses to burst stimulation.

### High-frequency burst spiking in dopaminergic axons is followed by refractory inhibition

We showed that the Ca2+ signal evoked by high-intensity nAChR activation results are enhanced following application of DHβE, suggesting that nAChRs can also play a role in reducing dopaminergic axon excitability. To test this, we recorded Ca2+ signals while stimulating high-frequency bursts (5 stimulations at 50 Hz) in the striatum (Figure 5A). We differentiated the signals to separately analyze the direct and nAChR components. As reported earlier (Figure 4E), the differentiated Ca2+ signal immediately following P1 had a biphasic waveform, with an early initial direct DA and a late nAChR-mediated component. For pulses P2-5, however, there were no peaks observed suggesting that stimulation during the period following P1 was ineffective in evoking axonal Ca2+ signals (differentiated peak Ca2+, P1: 0.127 +/- 0.03; P2: -0.003 +/- 0.003; P3: 0.014 +/- 0.002; P4: 0.019 +/- 0.003; P5: 0.023 +/- 0.005; n=16; Figure 5A and 5E).

**Figure 5.**
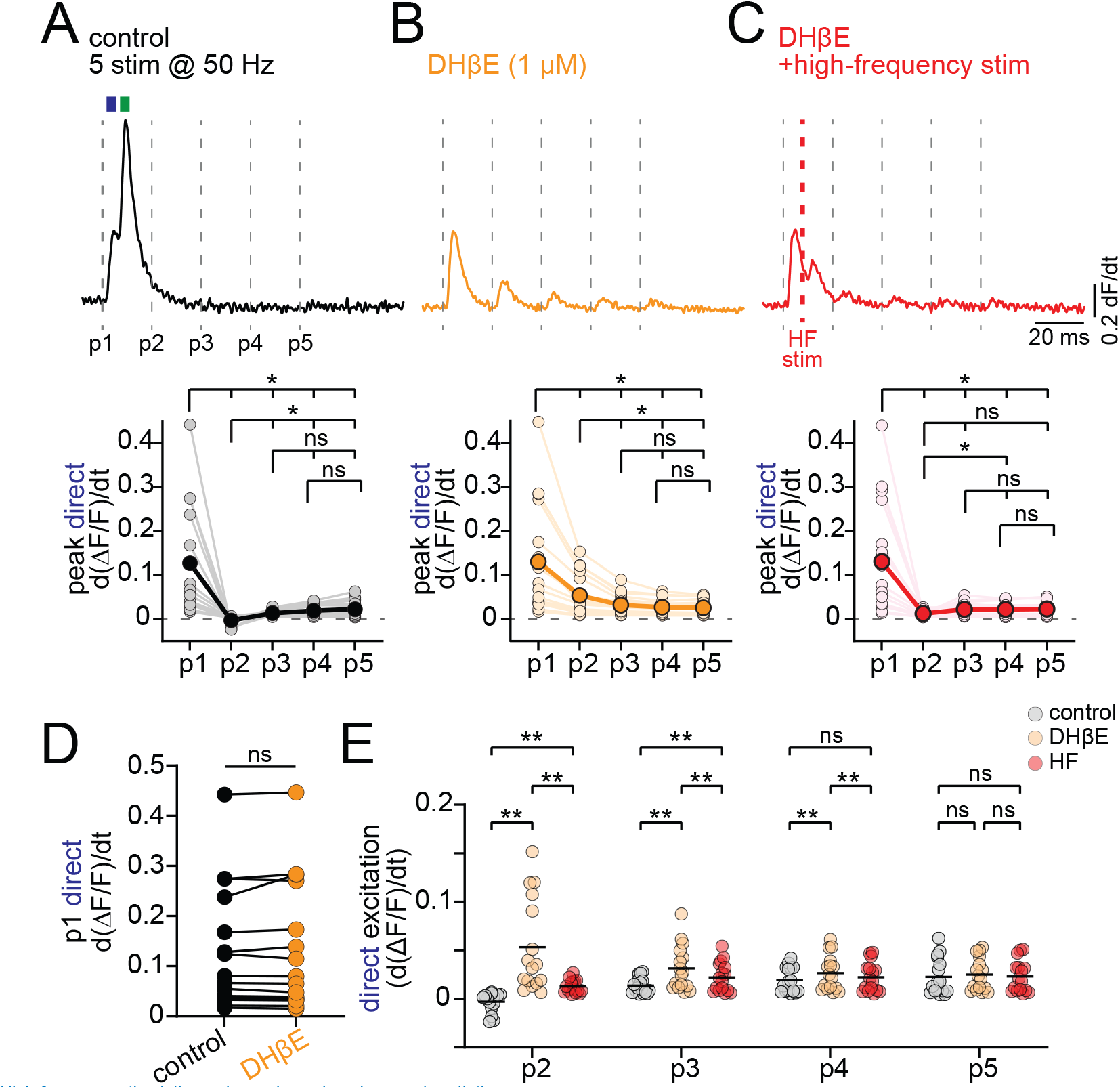
High-frequency stimulation reduces dopaminergic axonal excitation. **A.** (Top) Differentiated jGCaMP8f signal during 50 Hz DMS stimulation in control aCSF; dashed lines indicate individual pulses (P1–P5). Blue and green rectangles highlight direct and nAChR-mediated excitation, respectively. (Bottom) Group data for peak direct excitation across all 5 pulses. **B.** (Top) Example trace in DHβE; (Bottom) group data for peak direct excitation across 5 pulses. **C.** (Top) Example trace in DHβE with added electrical stimulation 6.8 ms after P1; (Bottom) group data for peak direct excitation across 5 pulses. **D.** Peak direct DAergic axon excitation (differentiated) for P1 in control and DHβE. **E.** Peak direct excitation for P2–P5 in control (grey), DHβE (orange), and DHβE with added high-frequency stimulation (red). *p<0.05, **p<0.01, ns > 0.05. Experiments performed in D2R, GABA-B, and mAChR antagonists.

Following DHβE block of nAChRs, stimulation during pulses P2-4 was significantly more effective in evoking Ca2+ signals, which then showed measurable peak amplitudes (avg. differentiated Ca2+ signal amplitude for control vs DHβE; P2: 0.056, q(15)=5.55, p=0.004; P3: 0.018, q(15)=5.79, p=0.003, P4: 0.007, q(15)=5.77, p=0.003; Figure 5B and 5E), arguing that the inhibition was mediated by nAChR signaling. Of note, there was no significant difference between the differentiated peak signals of control and DHβE by the pulse 5 of the train, which occurs 80 ms after the initial stimulation (mean difference ctrl vs DHβE P5: 0.002, q(15)=2.53, p=0.21; Figure 5E), suggesting the initial mechanism of inhibition is limited to about 80 ms in duration. As a control for the effect of inhibiting nAChRs on total axonal excitability, we also quantified the amplitude of the direct dopaminergic axon excitation following the first pulse, before the nAChR signaling occurs, and found there was no significant change by DHβE (ctrl=0.13, DHβE =0.13, q(15)=1.07, p=0.74; Figure 5D).

We hypothesized that if the inactivity period observed during pulse 2-5 reflects refractoriness caused by the nAChR-evoked high-frequency firing rather than a separate process, then it should be replicable in DHβE by using high-frequency stimulation to mimic the observed nAChR-dependent burst. To test this hypothesis, we modified the high-frequency burst stimulation protocol (HF stim) by inserting an additional pulse at 6.8 ms following pulse 1 (Figure 5C). With the extra pulse, the HF burst stimulation frequency following P1 was 147 Hz, which is comparable to the average axonal burst frequency shown in Figure 2. We found that the modified burst train significantly reduced the peak amplitude of differentiated Ca2+ signals for P2, P3 and P4 relative to the same signals recorded in DHβE (differentiated Ca2+ signal DHβE-HF; P2, q(15)=5.04, p=0.007; P3, q(15)=5.83, p=0.002; P4, q(15)=5.97, p=0.002; Figure 5E).

Thus, these data demonstrate that strong activation of nAChR results in a high-frequency burst of APs in dopaminergic axons that can lead to reduced axonal excitability for a period lasting 60-80 ms. Specifically, the period of inactivity likely reflects axon refractoriness that results from a high-frequency burst of AP firing caused by the activation of nAChRs.

## DISCUSSION

Here, we examined the mechanisms by which nAChR signaling both positively and negatively modulates the excitability of dopaminergic axons in the striatum. Calcium imaging experiments showed that nAChRs promote dopaminergic axon excitability at weak and moderate stimulation strengths. Using direct recordings from dopaminergic axons, we found that synchronous nAChR activation can evoke high-frequency bursts of APs in the striatum, which supports the role of nAChRs as excitatory receptors in dopaminergic axons. We also found that the axonal bursting evoked by strong stimulation is followed by a period of reduced axonal excitability. This refractory period in striatal dopaminergic axons is a consequence of the preceding AP burst firing and reduces further AP initiation in the distal axon for up to 80 ms. Our findings reveal a predominantly excitatory role for CINs over dopaminergic axonal excitability across most stimulation intensities, while also demonstrating that strong excitation can trigger high-frequency AP bursting, followed by a period of reduced axonal excitability. Thus, we explain how nAChRs can enhance and suppress dopaminergic axon excitability and striatal dopamine transmission, thereby accounting for the apparent low-pass filtering of striatal dopamine release by nAChR activity. Finally, our data further support the idea that dopamine axons can act as dynamic integrators of local network activity rather than simple signal propagators.

### nAChR subthreshold signaling is mainly excitatory

The evidence that nAChR signaling in dopaminergic axons is excitatory is supported by three main observations. First, nAChR signaling in dopaminergic axons produces robust AP firing, including a novel form of bursting we describe above. Second, local puff application of ACh elicits a subthreshold nAChR depolarization that increases the likelihood of AP initiation. Finally, except for under the strongest intensity stimulations, nAChR activity enhances calcium signaling in dopaminergic axons during both single and burst protocols. These results indicate that nAChRs enhance axonal excitability.

Previous work proposed that activation of axonal nAChRs can suppress DA release by depolarizing the axon and inactivating Na+ channels (Zhang et al., 2025), similar to the effects of axonal GABA-A receptors (Kramer et al. 2020). Although both are depolarizing, our observations here demonstrate that nAChRs and GABA-A receptors have opposite effects with activation of GABA-A receptors resulting in inhibition while nAChRs have mainly excitatory effects on axonal excitability. This divergence in function likely stems from the different characteristics of the nicotinic versus GABAa receptors. Our prior work showed that axonal GABAa receptors suppress axonal excitability due to a combination of depolarization and shunting inhibition. This is consistent with foundational work on presynaptic inhibition, which demonstrates that GABAa-mediated inhibition in axons is primarily due to shunting inhibition (Cattaert & El Manira, 1999; Eccles et al., 1961). By contrast, our experiments in Figure 3 show that activation of nAChRs produces little to no shunt. Furthermore, the reversal potential significantly more depolarized in nAChR relatively to GABA-A receptors (Erev, -6 mV vs -56 mV), which means that the driving force at subthreshold voltages will be significantly larger for nAChRs. Thus, the effect of nAChR activation is to depolarize the axonal membrane potential, bringing it closer to the spike threshold and thereby enhancing axonal excitability. Interestingly, axonal GABAa receptors are not always inhibitory. GABAa-mediated depolarizations in the cerebellum potentiate synaptic transmission (Pugh & Jahr, 2011), and in certain axons of the spinal cord GABAa activation may enhances AP propagation past branch points (Hari et al., 2022). These divergent functional outcomes of an axonal depolarization highlight the complex nature of modulating excitability in the axonal compartment.

While subthreshold depolarization enhances the likelihood of generating local axonal spikes, prolonged depolarization can also reduce axonal excitability affecting propagating action potentials that are traveling from the soma to the terminals. We found that activation of both GABAa and nACh receptors reduced AP height as a result of depolarization and Na+ channel inactivation. The amplitude of the subthreshold depolarization will determine whether it results in enhancement or suppression of axonal excitability. The excitatory nAChR-mediated depolarizations following puff application of ACh shown in Figure 3 were comparable in amplitude to spontaneous axonal EPSPs that we previously reported in DA terminals (Kramer et al., 2022). However, the depolarizations observed during the stimulated burst envelope in the striatum were larger in amplitude (5-10 mV), which would be expected to further reduce AP height, consistent with our observations (see Figure 3). This large subthreshold depolarization may reduce AP peak enough to suppress neurotransmitter release synergistically with the burst-mediated refractory period, contributing to nAChR-mediated suppression of DA release reported in vivo experiments (Zhang et al., 2025).

### nAChRs do not act as low-pass filters for DA release

In past work, nAChRs were proposed to act as a low-pass filter on DA release, enabling DA transmission at low frequencies while inhibiting DA release from high-frequency bursting at rates above 25 Hz (Rice & Cragg, 2004; Zhang & Sulzer, 2004). The cellular mechanism mediating this filtering has remained largely unresolved. Here, we show that nAChR activation does not act as a low-pass filter but rather can evoke high-frequency spiking, with instantaneous firing rates of ∼120 spikes per second. However, this period of strong excitation and nAChR-mediated bursting occurs only during the onset of the nAChR depolarization, within the first 20 ms of nAChR signaling. Evidence for this comes from both direct voltage recordings and from population-level calcium derivatives. Following this short period of excitation, there is a longer period of refractory inhibition that is a consequence of the intense excitation. Because we can mimic the nAChR refractory inhibition in the presence of nAChR antagonists with only high-frequency electrical stimulations (see Figure 5), we propose this inhibitory mechanism is not unique to nAChR signaling. Any period of strong excitation in dopaminergic axons would be expected to produce an ensuing period of inhibition, following the model of refractory inhibition first described by Hodgkin and Huxley (Hodgkin & Huxley, 1952). When evoking a train of stimulations at 50-100 Hz, many of the stimulation pulses will fall within this window of refractory inhibition. Since the refractory inhibition is mediated by the nAChR-dependent spiking, it is abolished by nAChR antagonists. Thus, it would appear that nAChRs act as low-pass filters when measuring DA release while evoking trains of stimulation. Of note, we found that optogenetic activation of CINs produced AP trains in dopaminergic axons with significantly more spikes than we saw with electrical stimulation. Thus, optogenetic activation may bias experimental results towards a strong stimulation outcome that includes prolonged refractory periods of inhibition.

### Axonal bursts result in an activity-dependent reduction in excitability

The observation that high-frequency firing results in a reduction in excitability has long been known in the cell bodies of dopaminergic neurons (Shepard & Bunney, 1991), but has not been observed in the axons. Here, we show by evoking AP firing in dopaminergic axons that the excitability of the axonal membrane also decreases over the duration of a high-frequency evoked spike train. We observed that stimulation of high-frequency activity progressively reduces the probability of axon AP initiation and delays the time to AP onset during a 10-pulse burst at 50 Hz. The mechanism of this adaptation is likely to be prolonged Na+ channel inactivation, which could occur through a slow-inactivation gate (Do & Bean, 2003) that is suggested to be present in the soma and dendrites of DANs (Ding et al., 2011). This period of refractory inhibition also resembles depolarization block, reduced excitability as a consequence of intense activation (Blythe et al., 2009; Lammel et al., 2008; Richards et al., 1997; Tucker et al., 2012). It is also possible that axonal K+ channels are activated over the duration of a burst that then decreases excitability. One such channel is KCNQ4, which has been shown to modulate DA release when activated in striatal sections (Jensen et al., 2011), suggesting these channels are expressed in dopaminergic axons.

### Action potential bursting initiated in the axons of dopaminergic neurons is a new functional output signal

Dopaminergic somata of the ventral midbrain have two main firing modes, tonic and bursting. Tonic firing is linked to movement behaviors and is thought to set a baseline level of DA release. By contrast, AP bursting of midbrain DANs is a phasic signal that drives DA-dependent learning in response to salient sensory stimuli (Hart et al., 2014; Kutlu et al., 2021; Schultz et al., 1997). The difference between the tonic DA signal and the burst-mediated DA signal may convey information about the sensory experience. Unlike bursting in the soma, which typically occurs around 20 Hz, the burst firing we observed in dopaminergic axons occurs at rapid frequencies of up to 200 Hz (Grace & Bunney, 1983; Otomo et al., 2020). Therefore, the ultra-fast burst of APs evoked by CIN input onto dopaminergic axons may convey different information into the striatum that is distinct from somatic bursting. However, whether CINs evoke bursting in DAergic axons in vivo is yet to be determined.

We report that the nAChR-evoked bursting occurs at a high-frequency of 180 Hz which is within the detection range of direct patch clamp recordings with submillisecond resolution. Using the fast kinetics of genetically-encodable calcium indicators (GCaMP8f) coupled with photometry, our studies enable separate analysis of direct axonal calcium signals and nAChR generate signals that are separated by 8-9 ms (Brill-Weil et al, 2025). Detecting the rapid and high-frequency DA release would require a high image acquisition rate combined with a fast DA sensor capable of detecting DA release within 20 ms. A recent study showed that nAChR-evoked dopamine release is detectable in vivo when using the DA sensor dLight1.2 (∼10 ms on rate) combined with a high frame rate (Flink et al., 2025). Thus, further studies are required using rapid detection methods in order to determine the behavioral conditions under which CINs produce single action potential-, or burst-, mediated DA release.

## METHOD

### Mice

All animal handling and procedures were approved by the animal care and use committee (ACUC) for the National Institute of Neurological Disorders and Stroke (NINDS) at the National Institutes of Health. Mice of both sexes were used throughout the study. Mice that underwent viral injections were injected at postnatal day 21 or older and were used for ex vivo imaging 1-4 weeks after injection. DAT-Cre (RRID:IMSR_JAX:006660) and Ai9 (RRID:IMSR_JAX:007909) strains were used.

### Viral injections

All stereotaxic injections were conducted on a Stoelting QSI (Cat#53311). Mice were maintained under anesthesia for the duration of the injection and allowed to recover from anesthesia on a warmed pad. The viruses used for this study were jGCaMP8f (AAV9-pGP-syn-FLEX-jGCaMP8f-WPRE titer: > 1x 1013, Addgene #162379) and cre-dependent tdTomato (AAV9-CAG-FLEX-tdTomato titer: > 1x 1012, plasmid: Addgene #28306). Viral aliquots were injected (0.1-0.5 μl) bilaterally into the SNc (X: ± 1.9 Y: -0.5 Z: -3.9) via a Hamilton syringe. At the end of the injection, the needle was raised at a rate of 0.1 to 0.2 mm per minute for 10 minutes before being removed.

### Tissue sectioning

Experiments were performed on male and female adult mice of at least 6 weeks in age. Mice were anesthetized with isoflurane, decapitated, and brains rapidly extracted. Horizontal sections were cut at 330 μm thickness on a vibratome while immersed in warmed, modified, slicing ACSF containing (in mM) 198 glycerol, 2.5 KCl, 1.2 NaH2PO4, 20 HEPES, 25 NaHCO3, 10 glucose, 10 MgCl2, 0.5 CaCl2. Cut sections were promptly removed from the slicing chamber and incubated for 30-60 minutes in a heated (34°C) chamber with holding solution containing (in mM) 92 NaCl, 30 NaHCO3, 1.2 NaH2PO4, 2.5 KCl, 35 glucose, 20 HEPES, 2 MgCl2, 2 CaCl2, 5 Na-ascorbate, 3 Na-pyruvate, and 2 thiourea. Slices were then stored at room temperature and used 30 min to 8 hours later. Following incubation, slices were moved to a heated (33–35°C) recording chamber that was continuously perfused with recording ACSF (in mM): 125 NaCl, 25 NaHCO3, 1.25 NaH2PO4, 3.5 KCl, 10 glucose, 1 MgCl2, 2 CaCl2. In experiments where there were drugs in the circulating ACSF, the slices were incubated in the recording solution for at least 15 minutes before recording began.

### Electrophysiology and functional imaging

For imaging experiments, a white light LED (Thorlabs; SOLIS-3C) was used in combination with a EGFP (Chroma; 49002) filter set to visualize DAergic axons infected with jGCaMP8f. For visualizing the jGCaMP8f signals, a photodiode (New Focus) was mounted on the top port of the Olympus BX-51WI. Signals were using a Digidata 1550 (Molecular Devices) sampled at 50kHz. All electrical stimulation was delivered with tungsten bipolar electrodes (250 μm tip separation, MicroProbes) placed 100-200 μm from the imaging site in the DMS. Electrical stimulation was given using an Isoflex (A.M.P.I.) with amplitudes ranging from 0.1 to 3 V. For stimulation-response curves, a minimal stimulation intensity was experimentally determined for each replicate by reducing the stimulation intensity until there was no observable response. Then, the intensity was increased until a minimal response emerged, which was set as the “minimal” intensity (1). Stimulations were then increased as multiples of this normalized minimal intensity (3x, 10x, and 30x). Experiments in “iso. aCSF” were in the presence of GABAb (CGP-55845, 300 nM), mAChR (atropine, 100 nM), and D2R (sulpiride, 300 nM) inhibitors.

Perforated-patch recordings from striatal TdTomato+ axons were made using borosilicate pipettes (5-9 MΩ) filled with internal solution containing (in mM) 135 KCl, 10 NaCl, 2 MgCl2, 10 HEPES, 0.5 EGTA, 0.1 CaCl2, adjusted to a pH value of 7.43 with KOH, 278 mOsm. Pipette tips were back-filled with ∼1 μL of clean internal. Pipettes were then filled with internal containing between 80 and 100 μg/mL gramicidin. Patch integrity was monitored by the addition of Alexa-488 to the gramicidin-containing internal. Whole-cell recordings from the TdTomato+ axons in the medial forebrain bundle were made using borosilicate pipettes (4-7 MΩ) filled with internal solution containing (in mM) 122 KMeSO3, 9 NaCl, 1.8 MgCl2, 4 Mg-ATP, 0.3 Na-GTP, 14 phosphocreatine, 9 HEPES, 0.45 EGTA, 0.09 CaCl2, adjusted to a pH value of 7.35 with KOH. All recordings were made with a MultiClamp 700B (Molecular Devices).

Pressure ejection of ACh was performed using borosilicate pipettes (2-4 MΩ). ACh (300 μM) was added to a modified external solution containing (in mM): 125 NaCl, 25 NaHCO3, 1.25 NaH2PO4, 3.5 KCl, 10 HEPES, 0.01 Alexa 488, final osmolarity 280–290 mOsm. This puffing solution was then spin filtered, loaded into a glass pipette, and lowered to within 30–50 μm of the axon using a micro-manipulator. The puffing solution was applied onto the axon with a short pressure ejection (100-250 msec in duration) using a PV 820 Pneumatic PicoPump (WPI).

The starting amplitude of the current injection protocol in Figure 4 was experimentally tuned for each axon to be close to rheobase (*I*_Rh_), and increased by 5 pA for each subsequent trial (Figure 4B). 10 current steps were performed with a 20 ms inter-stimulus interval. The protocol was continued until a 100% success rate of spiking was achieved. For analysis, only sweeps prior to the 100% successes of AP initiation were considered.

### Quantification and statisitcs

Analysis was conducted in Igor Pro (Wavemetrics) and statistical tests were performed in Prism 9 (GraphPad) and Igor Pro. Parametric data are reported as the mean while non-parametric are reported as the median. Error bars for non-parametric data are shown as standard deviation (s.d.), while those for parametric data are standard error of the mean (s.e.m.). Data in box-and-whisker plots (non-parametric) are showns as: the box line is at the median, the hinges of the box extend from the 25th to 75th percentiles, and the whiskers extend from the minimum to maximum in the dataset. T-tests were used for two-group comparisons, and ANOVA tests were used when comparing more than two groups followed by a Bonferroni post-hoc test for analysis of multiple comparisons. Cumulative distributions were compared with a Kolmogorov-Smirnov test. Statistics are shown t(df)=t_val where df indicates the degrees of freedom and Y indicates the calculated test statistic.

#### Action potential analysis

To quantify APs, the derivative was taken of each trace in the recording and the stimulation artifacts were blanked. The derivative traces were analyzed in Igor Pro using NeuroMatic (Rothman & Silver, 2018). A cutoff value was set to identify AP derivatives that were at least 2 standard deviations above the baseline noise. Derivatives were then manually confirmed to be APs. The AP success rate in Figure 4 was calculated across axons for each stimulation pulse (1 through 10) in control (black) and during ACh ejection (green).

#### Discrimination between direct and nicotinic components

For discrimination between the direct and nicotinic components of the evoked jGCaMP8f signal, the average waveform of at least 2 evoked Ca2+ events were taken under control conditions. The “direct component” time window was defined as the time between the onset (5% of peak) of the first peak and the onset of the second peak. This resulted in a region that was a 5.6 ms window beginning 1.1 ms after the stimulus. The analysis window for the nicotinic component was defined as a 20 ms window immediately following the direct component. This analysis is shown in Figure 4. The analysis starts with a raw jGCaMP8f signal, which is first differentiated. The application of DHβE defines the direct differentiated signal in DAergic axons. The direct differentiated signal is then subtracted from the control signal to produce an isolated nAChR-mediated differentiated signal.

#### Amplitude quantification of direct and nicotinic components

In order to quantify changes to the direct component of evoked jGCaMP8f signals, the first derivative of the raw signal was low pass filtered. The maximum value within the direct component analysis window was determined for every trace in each slice to create a time series. To quantify the nicotinic component for each slice, the average first derivative trace in DHβE was subtracted from each individual first derivative wave of the recording. This resulted in first derivative waves that contained only the nicotinic contribution to the jGCaMP8s signal. To make the analysis robust to noise and account for any changes in the direct component amplitude, each individual trace was then baselined to the second half (2.8 msec) of the direct component window. The peak value was taken for every wave within the analysis window for the nicotinic component, as defined above. To account for baseline noise in the first derivative photometry traces, the average noise was determined for each slice using a 5.6 msec window immediately before the stimulus. This average value from all waves in each slice was subtracted from the quantification of the peak nicotinic signal. Finally, both the quantifications of direct and adjusted nicotinic components for each slice were normalized to control conditions and averaged across slices. Quantification of drug effects was done by averaging the final 5 minutes of the control or wash-in periods.

## Acknowledgements

This work was supported by NINDS Intramural Research Program grant NS003135 to Z.M.K. and by startup Funds from the Department of MCDB, University of Michigan (PFK). Funding for this work was also provided by Aligning Science Across Parkinson’s (ASAP-020529) to Z.M.K. through the Michael J. Fox Foundation for Parkinson’s Research (MJFF). The contributions of the NIH author(s) were made as part of their official duties as NIH federal employees, are in compliance with agency policy requirements, and are considered Works of the United States Government. However, the findings and conclusions presented in this paper are those of the author(s) and do not necessarily reflect the views of the NIH or the U.S. Department of Health and Human Services.

## Author Contributions

PFK, ZMK conceived the project. PFK, AY, FC, RZ, ZMK designed and performed experiments. PFK, AY, FC, ZMK analyzed the data. PFK wrote the first draft of the manuscript and PFK and ZMK edited the manuscript to produce the final version.

---

**Supplemental Figure 1.**
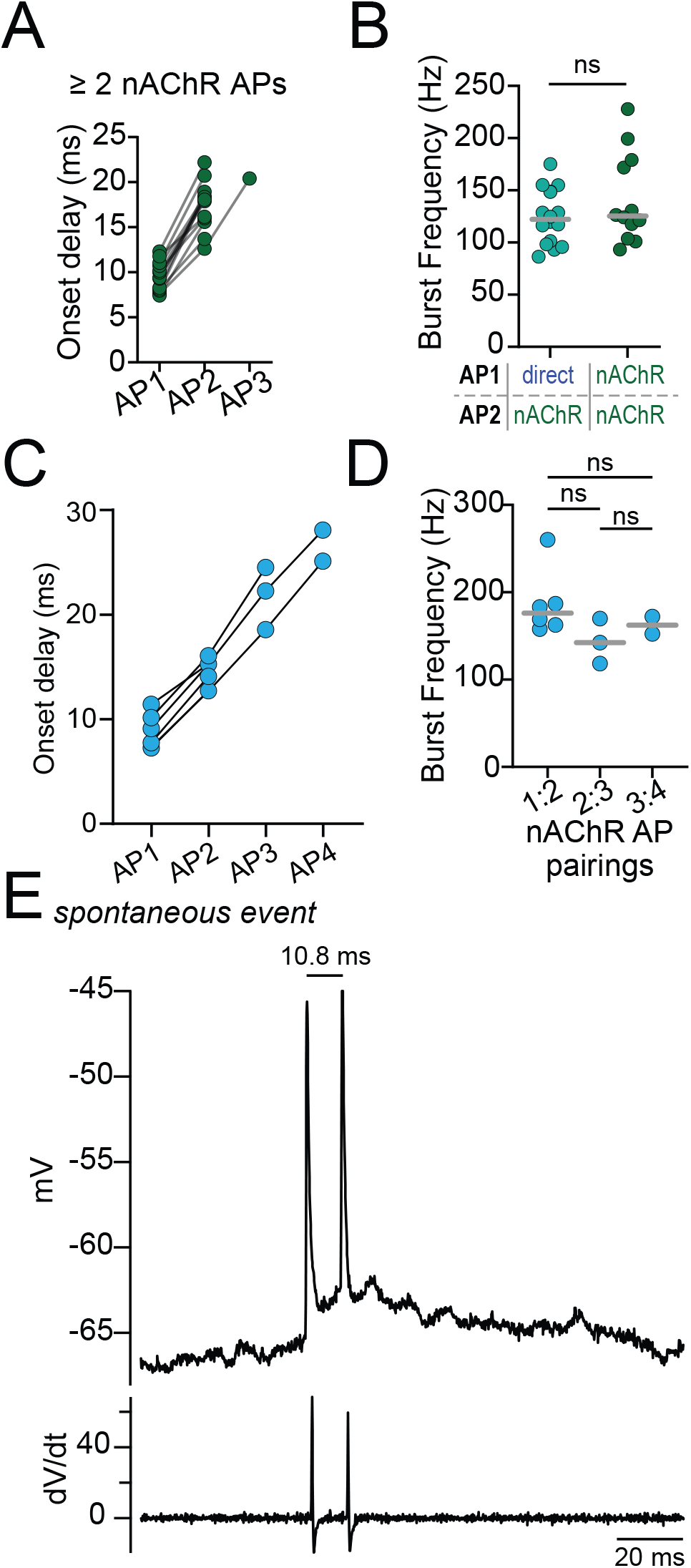
Further characterization of bursting in dopaminergic axons. **A.** Action potential onset delay following a single electrical stimulation for nAChR-mediated spikes in records with > 2 events. **B.** No significant difference in the instantaneous frequencies between pairs of APs that combine a direct and nAChR-mediated spike, or two nAChR-mediated spikes. **C.** Action potential onset delay following a single LED light pulse for nAChR-mediated spikes in records with > 2 events. **D.** No significant difference in the instantaneous frequency between pairs of APs throughout the LED-evoked burst train. **E.** Example recording of a spontaneously generated high-frequency burst of APs recorded in a DAergic axon on the DMS.

**Supplemental Figure 2.**
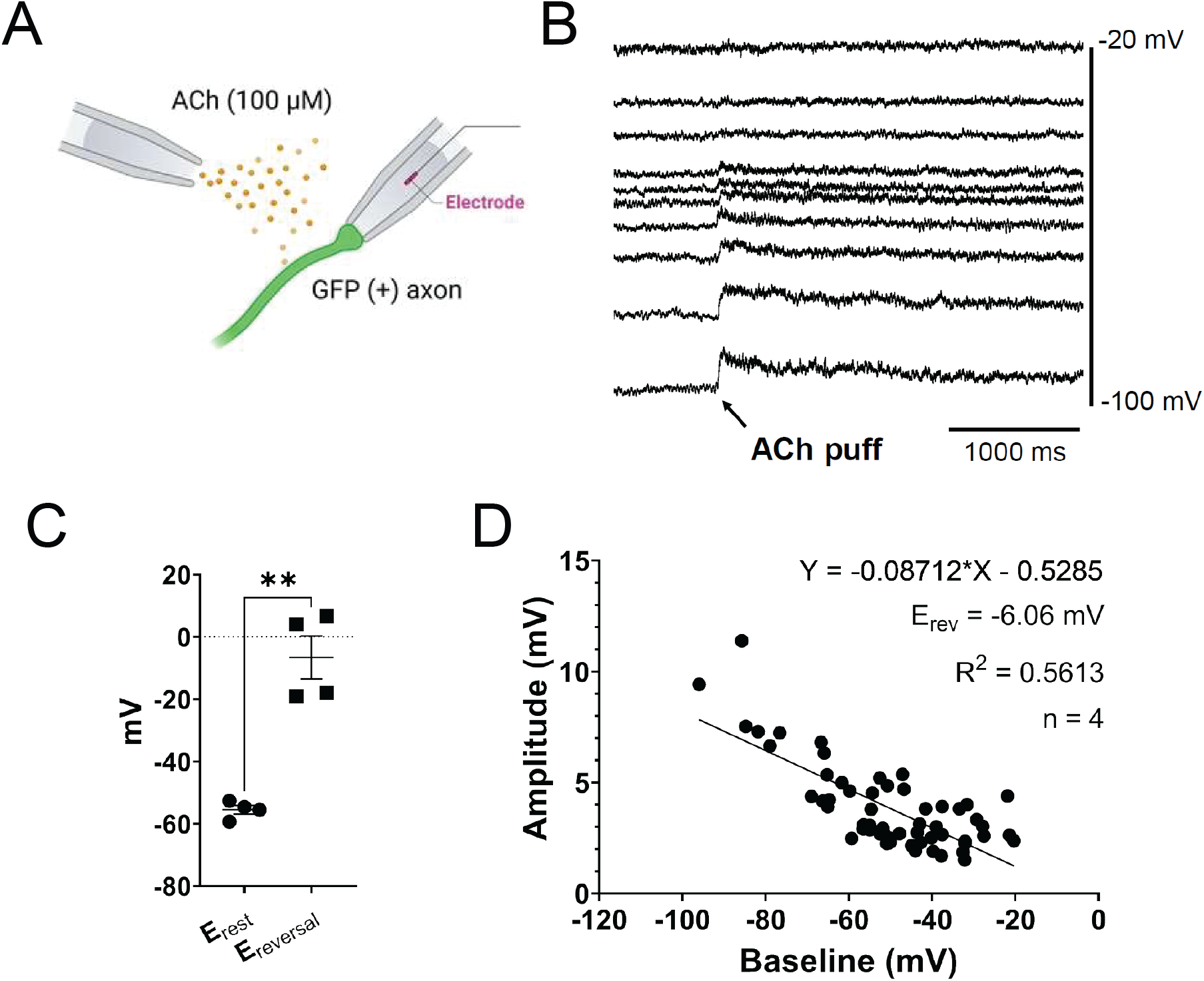
nAChR potentials reverse near 0 mV in dopaminergic axons. **A.** Experimental setup: DAergic axons in the MFB were recorded using perforated patch recordings from MFB axons; nAChRs activated by local ACh ejection. **B.** Example recording showing an ACh-mediated depolarization at different resting membrane potentials. **C.** Combined data showing a significantly depolarized reversal potential for nAChRs relative to the axonal resting membrane potential. **D.** Combined data showing the calculation of a reversal potential using linear estimation. ** p<0.01

